# Interaction of taste and place coding in the hippocampus

**DOI:** 10.1101/431353

**Authors:** Linnea E. Herzog, Leila May Pascual, Seneca J. Scott, Elon R. Mathieson, Donald B. Katz, Shantanu P. Jadhav

## Abstract

An animal’s survival depends on finding food, and the memory of food and contexts are often linked. Given that the hippocampus is required for spatial and contextual memory, it is reasonable to expect related coding of space and food stimuli in hippocampal neurons. However, relatively little is known about how the hippocampus responds to tastes, the most central sensory property of food. In this study, we examined the taste-evoked responses and spatial firing properties of single units in the dorsal CA1 hippocampal region as male rats received a battery of taste stimuli differing in both chemical composition and palatability within a specific spatial context. We identified a subset of hippocampal neurons that responded to tastes, some of which were place cells. These taste and place responses had a distinct interaction: taste-responsive cells tended to have less spatially specific firing fields, and place cells only responded to tastes delivered inside their place field. Like neurons in the amygdala and lateral hypothalamus, hippocampal neurons discriminated between tastes predominantly on the basis of palatability, with taste-selectivity emerging concurrently with palatability-relatedness; these responses did not reflect movement or arousal. However, hippocampal taste responses emerged several hundred msec later than responses in other parts of the taste system, suggesting that the hippocampus does not influence real-time taste decisions, instead associating the hedonic value of tastes with a particular context. This incorporation of taste responses into existing hippocampal maps could be one way that animals use past experience to locate food sources.

**Significance statement:** Finding food is essential for animals’ survival, and taste and context memory are often linked. While hippocampal responses to space and contexts have been well characterized, little is known about how the hippocampus responds to tastes. Here, we identified a subset of hippocampal neurons that discriminated between tastes based on palatability. Cells with stronger taste responses typically had weaker spatial responses, and taste responses were confined to place cells’ firing fields. Hippocampal taste responses emerged later than in other parts of the taste system, suggesting that the hippocampus does not influence taste decisions, but rather, associates the hedonic value of tastes consumed within a particular context. This could be one way that animals use past experience to locate food sources.

## Introduction

The hippocampus is essential for spatial learning and memory, and is thought to provide a cognitive map of animals’ experience. The central data for this view come from studies of place cells that respond to specific locations as animals explore their environments (O’Keefe and Nadel, 1978; Moser et al., 2008).

Given that one of the most obvious uses for such a mental map is to aid in the finding of food, it is surprising how little is known about how the hippocampus processes taste, the most central sensory property of food. It is reasonable to expect that taste information reaches the hippocampus; although not traditionally considered to be part of the taste system, anatomical studies show that the hippocampus receives projections, either directly or indirectly through the entorhinal cortex, from several brain regions in which taste information is processed, including the gustatory cortex (GC), orbitofrontal cortex, and amygdala (Suzuki and Amaral, 1994; von Bohlen und Halbach and Albrecht, 2002). Functional imaging studies in humans also indicate that the hippocampal formation is active during taste ingestion and discrimination (Zald et al., 1998; Haase et al., 2009; Spetter et al., 2010), and rodent lesion studies suggest that the hippocampus plays a role in taste learning (Reilly et al., 1993; Stone et al., 2005; Chinnakkaruppan et al., 2014).

A great deal is known about taste responses in other parts of the taste system: in cortex, these responses evolve dynamically, reflecting taste presence, identity and palatability in distinct epochs preceding the decision to consume or expel a given taste (Katz et al., 2001; Sadacca et al., 2012; Sadacca et al., 2016). However, it remains unclear if or how hippocampal taste responses co-exist and interact with representations of space. While hippocampal neurons are known to respond to tastes in a context-dependent manner (Ho et al., 2011), no studies to date have directly measured single unit responses to tastes in the hippocampus alongside spatial firing properties, or examined the dynamics of these responses.

Hippocampal place cells are certainly capable of encoding non-spatial information such as odors (Wood et al., 1999), visual cues (Fried et al., 1997), textures (Shapiro et al., 1997), tones (Moita et al., 2003) and time (Kraus et al., 2013). Place cells typically respond to these stimuli by modulating their firing rate (“rate remapping,” Leutgeb et al., 2004; Allen et al., 2012) or firing location (“global remapping,” Leutgeb et al., 2005; Fyhn et al., 2007). A new cognitive map can also be formed based on the parameters of a behaviorally relevant non-spatial stimulus (Kraus et al., 2013; Aronov et al., 2017). The difficulty inherent in dissociating spatial from non-spatial influences in behaving rodents (O’Keefe, 1999), however, has led some researchers to propose that seeming responses to non-spatial stimuli may simply reflect arousal triggered by the onset of the stimulus, rather than true sensory responses (Shan et al., 2016). To establish that non-spatial responses are genuine, it is necessary to show that spatially tuned neurons can discriminate between sensory stimuli.

Here, we did just this, recording single-unit activity in the dorsal CA1 region of awake rats while exposing them to four taste solutions. We identified subsets of place cells and interneurons that discriminated between tastes based predominantly on palatability; this pattern was consistent with those observed in basolateral amygdala (BLA; Fontanini et al., 2009) and lateral hypothalamus (LH; Li et al., 2013), although hippocampal taste dynamics evolved much more slowly. Neurons classified as taste-responsive place cells responded exclusively to tastes delivered within their place field, and tended to have lower spatial selectivity than non-taste-responsive place cells. Together, these results establish that hippocampal responses to sensory stimuli do not simply reflect changes in arousal state, and can encode sensory parameters relevant for behavior. Further, they suggest that hippocampal taste responses may be used to form value-related associations between tastes and contexts, which can facilitate using past experience to locate food sources.

## Materials and Methods

### Animals and surgery

Five adult (450-550 g) male Long-Evans rats (Charles River Laboratories, RRID: RGD_2308852) were used as subjects in this study. Rats were kept on a 12 h light/dark cycle, with all sessions taking place around the same time during the light period. All surgical and experimental procedures were conducted in accordance with the National Institutes of Health guidelines and approved by the Brandeis University Institutional Animal Care and Use Committee.

After several weeks of habituation to daily handling, animals were chronically implanted with a microdrive array consisting of 25-30 independently moveable tetrodes in the right dorsal hippocampal region CA1 (−3.6 mm AP, 2.2 mm ML), and an intra oral cannula (IOC). Each IOC consisted of a polyethylene tube inserted beneath the temporalis muscle and terminating anterolateral to the first maxillary molar, allowing for the precise delivery of taste solutions onto the rat’s tongue (Grill and Norgren, 1978; Travers and Norgren, 1986; Katz et al., 2001).

Following recovery from the implantation surgery (~7-8 days), rats were water-deprived to 85-90% of their *ad libitum* weight to ensure taste consumption during the recording sessions. At ~14 d after implantation, animals were habituated for at least 3 days to the behavioral chamber, sleep box, and the delivery of taste solutions through the IOC; place cells were expected to be stable after habituation (Thompson and Best, 1990; Agnihotri et al., 2004). Following habituation, we performed daily recording sessions in which rats were exposed to pseudo-randomized sequences of four standard taste stimuli (see **Figure 1A, 1B** and *Passive taste administration paradigm*). Following the conclusion of experiments, we made electrolytic lesions through each electrode tip to mark recording locations. Brains were sectioned into 50 µm slices and stained with cresyl violet to confirm electrode placement in the hippocampal cell layer (see **Figure 1C**).

**Figure 1.**
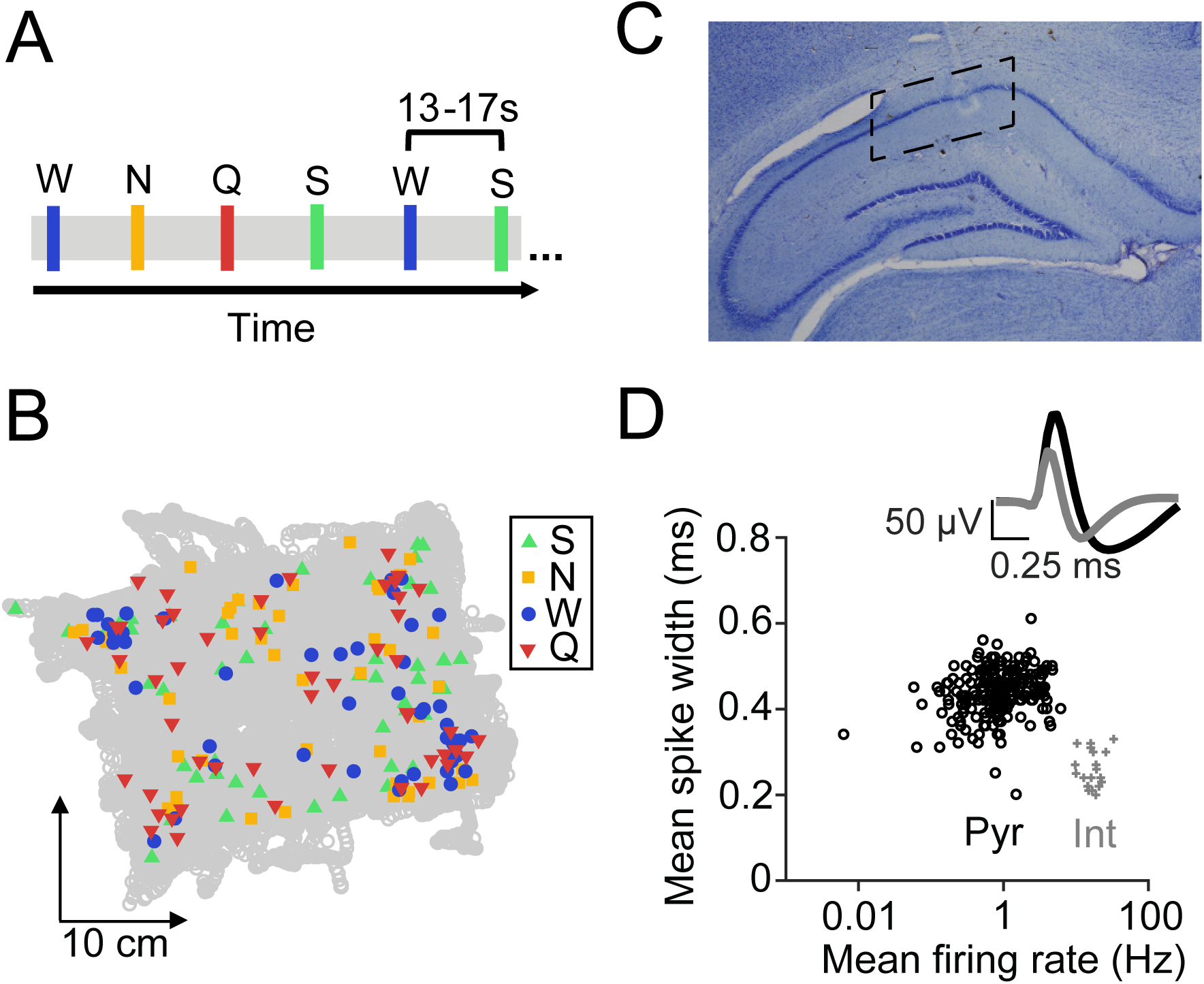
Experimental design and electrophysiology. ***A***, A portion of the timeline of an example taste delivery experiment. Colored bars indicate individual deliveries of taste stimuli: green (S, 4 mM saccharin), yellow (N, 100 mM sodium chloride), blue (W, distilled water) and red (Q, 5 mM quinine hydrochloride). Taste deliveries occurred at a randomized timing of 13-17 s, with the taste identity randomized for each trial. ***B***, Example session showing all 200 taste delivery locations (colored symbols) overlaid on top of the rat’s position in the behavioral chamber (gray circles) during one recording experiment. ***C***, Histological verification of tetrode locations in intermediate dorsal CA1. Dotted lines indicate the extent of recording sites across all five animals. ***D***, Classification of putative interneurons (Int, gray crosses) from pyramidal cells (Pyr, black circles) based on spike width (> 8.5 Hz) and firing rate (< 0.35 ms) parameters. The top panel depicts average waveform shapes for putative interneurons (gray) and pyramidal cells (black).

### Passive taste administration paradigm

Each recording session typically lasted between 2 and 3 hours, and consisted of three sessions in a ~30 x 35 x 40 cm Plexiglass behavioral chamber (J. Green, Charles River Maker Lab) interleaved with four 15-20 minute sleep sessions in a ~30 x 30 x 40 cm black box (rest box). The first and last sessions in the behavioral chamber consisted of 15-20 minute periods in which animals were habituated to the behavioral chamber in the absence of tastes. During the middle experimental session (depicted in **Figure 1A**), rats received a pseudo-randomized sequence of four standard taste stimuli [sweet: 4 mM saccharin (S); salty: 100 mM sodium chloride (N); neutral: distilled water (W); and bitter: 5 mM quinine hydrochloride (Q)] that varied in hedonic value and fell within the range of concentrations typically used by other groups (3-20 mM saccharin, 10-300 mM sodium chloride, and 1-10 mM quinine; for review, see Frank and Brown, 2003; Kobayakawa et al., 2005; Accolla and Carleton, 2008; Geran and Travers, 2009; Rosen et al., 2010; Chen et al., 2011; MacDonald et al., 2012; Li and Lemon, 2015; Sadacca et al., 2016). Taste solutions were delivered directly onto the tongue in ~40 µL aliquots via four polyamide tubes inserted into the IOC, with a separate tube for each solution to prevent the mixing of tastes. Rats received 50 pseudo-randomized repeats of each of the four taste stimuli, for a total of 200 taste deliveries. This number of deliveries allowed us to obtain sufficient numbers of in-field and out-of-field trials for each taste stimulus (see *In-field vs. out-of-field analysis*) before rats became satiated (Fontanini and Katz, 2005). This requisite number of trials per stimulus is much greater than what is typically used in studies of taste coding in awake rodents (10-30 trials, see Katz et al., 2001; Li et al., 2013; Baez-Santiago et al., 2016; Li et al., 2016), and therefore necessitated limiting the number of stimuli (e.g., sour taste) required for these comparisons (Moita et al., 2003). The interval between taste deliveries was randomized to 13-17 seconds, which is sufficiently long enough to prevent mixture effects or contamination by previous deliveries, even without rinses factored into the experimental paradigm, since awake rats have previously been shown to engage in what amounts to a constant saliva rinse (Fontanini and Katz, 2006). The total amount of fluid delivered in each ~50 minute taste administration period was 8 mL, after which animals had access to an additional 15-20 mL of water in their home cage.

### Electrophysiology

Electrophysiological recordings were conducted using a SpikeGadgets system (Tang et al., 2017). Spikes were sampled at 30 kHz and bandpass filtered between 600 Hz and 6 kHz. Local field potentials (LFPs) were sampled at 1.5 kHz and bandpass filtered between 0.5 and 400 Hz. During recording sessions, the animal’s position and speed were recorded using an overhead monochrome CCD camera (30 fps) and tracked by LEDs affixed to the headstage.

Over ~14 d following surgery, tetrodes were gradually advanced to the CA1 hippocampal cell layer, as identified by characteristic EEG patterns (sharp-wave ripples, or SWRs; theta0020 rhythm) as previously described (Jadhav et al., 2012; Jadhav et al., 2016; Tang et al., 2017). Tetrodes were readjusted after each day’s recordings. Each animal had one hippocampal reference tetrode in corpus callosum, which was also referenced to a ground screw installed in the skull overlying cerebellum.

Single units were isolated offline based on peak amplitude and principal components (Matclust, M.P. Karlsson). Only well-isolated units with stable waveforms that fired at least 100 spikes per session were included in our analysis. As is typical for studies of hippocampal CA1 neurons (Fox and Ranck, 1981; Jadhav et al., 2016; Tang et al., 2017), putative interneurons (*Int*) were identified on the basis of firing rate (> 8.5 Hz) and spike width (< 0.35 ms) parameters (**Figure 1D**). All other isolated units were classified as putative pyramidal cells (*Pyr*). We emphasize that these distinctions are putative, as definitive classifications of cell types cannot be determined from extracellular recordings alone. We isolated a total of 482 neurons from five rats, conducted across nine experiments. **Table 1** shows the distribution of cells across all five animals.

**Table 1.**
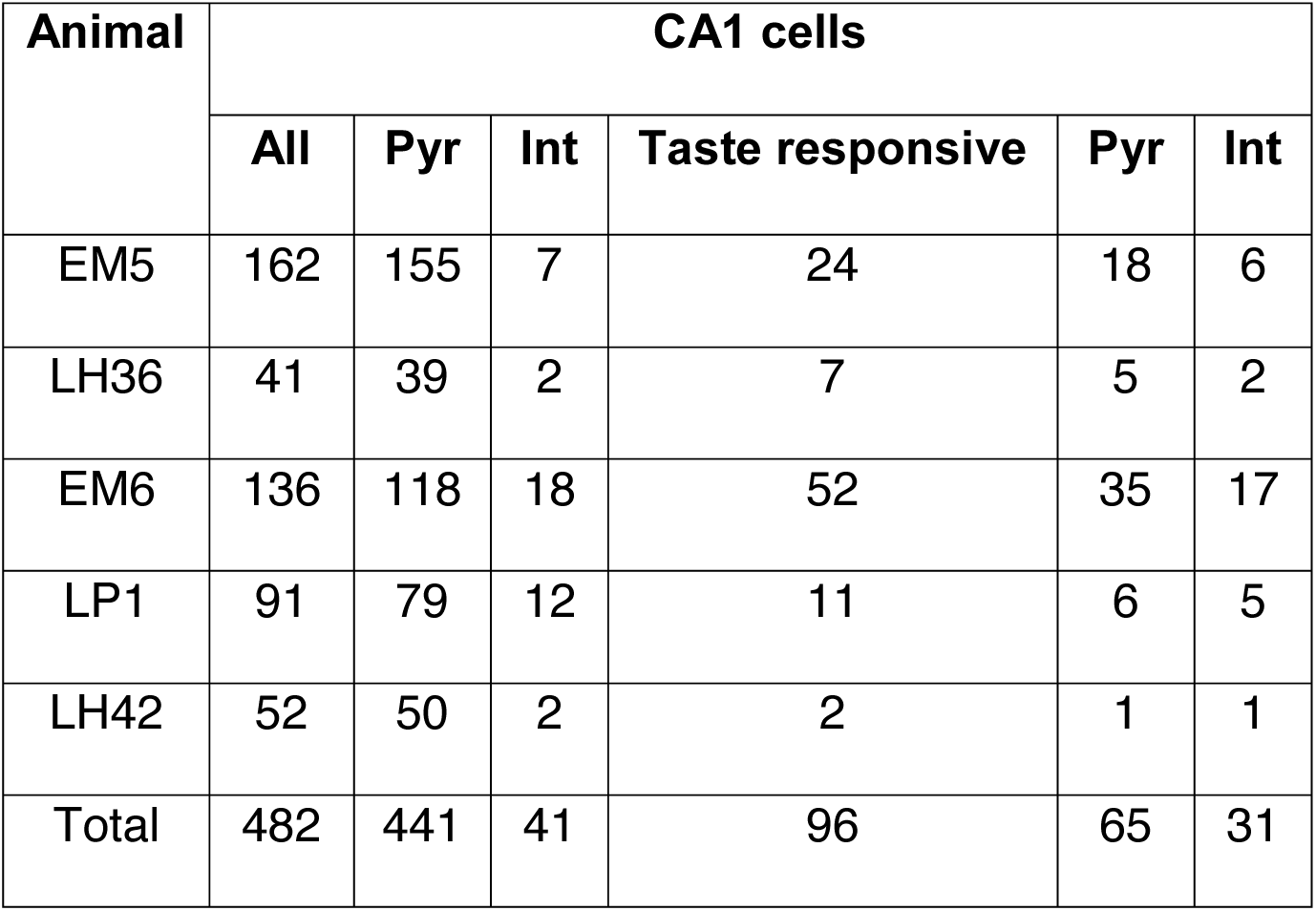
Cell distribution across animals. Summary of the number of taste-responsive and total CA1 cells recorded from each animal. Only the cells meeting the inclusion criteria (see Materials and Methods) are reported. Putative pyramidal cells (Pyr) and interneurons (Int) were identified on the basis of firing rate and spike width parameters. Neurons were classified as “taste-responsive” if they exhibited responses to taste presence, identity and/or palatability.

| Animal | CA1 cells |  |  |  |  |  |
| --- | --- | --- | --- | --- | --- | --- |
|  | All | Pyr | Int | Taste responsive | Pyr | Int |
| EM5 | 162 | 155 | 7 | 24 | 18 | 6 |
| LH36 | 41 | 39 | 2 | 7 | 5 | 2 |
| EM6 | 136 | 118 | 18 | 52 | 35 | 17 |
| LP1 | 91 | 79 | 12 | 11 | 6 | 5 |
| LH42 | 52 | 50 | 2 | 2 | 1 | 1 |
| Total | 482 | 441 | 41 | 96 | 65 | 31 |

### SWR detection

SWRs were detected as previously described (Jadhav et al., 2016; Tang et al., 2017) using the ripple-band (150-250 Hz) filtering of LFPs from multiple tetrodes. A Hilbert transform was used to determine the envelope of band-passed LFPs, and events that exceeded a threshold (mean + 3 SD) were detected. SWR events were defined as the times around initially detected events when the envelope exceeded the mean. SWR periods were excluded from place field analysis, similar to previous studies (Jadhav et al., 2016; Tang et al., 2017).

### Palatability/preference data

Palatability can be defined as the relative hedonic value of tastes; this characteristic is related, but not identical to reward, as palatability is a continuously-varying property of taste that is easily manipulated by experience (Stone et al., 2005; Katz and Sadacca, 2011; Chinnakkaruppan et al., 2014). Taste palatability was assessed using a brief-access task (BAT, Davis Rig Gustometer, Med Associates; for details, see Sadacca et al., 2016) in a separate cohort of adult male rats (n = 7) that underwent the same water restriction protocol as the rats used in the recording experiment. Consumption data were averaged across two testing days for each animal. The palatability rank order determined by the brief-access test (S > N > W > Q, see **Figure 6C**) matches what has been observed in numerous studies across a broad range of stimulus delivery methods and assessment techniques (Travers and Norgren, 1986; Breslin et al., 1992; Clarke and Ossenkopp, 1998; Fontanini and Katz, 2006; Sadacca et al., 2016).

### Experimental design and statistical analysis

#### Spatial maps

To characterize the spatial firing properties of neurons, two-dimensional occupancy-normalized firing rate maps (**Figures 2**, **4, 5**) were made using 0.5 cm square bins and smoothed with a 2D Gaussian (σ = 3 cm; Tang et al., 2017). Data from taste delivery (500 ms before to 2500 ms after) and SWR periods (see *SWR detection and modulation*) were excluded from spatial map analysis. Peak rates for each cell were defined as the maximum firing rate across all spatial bins in the spatial map. Each cell’s place field was defined as the largest cluster of neighboring spatial bins which had firing rates >= 20% of the peak rate; place field sizes were then calculated by multiplying the number of bins by the bin size (Brun et al., 2002). Unpaired t-tests were used to determine whether the mean firing field size differed significantly between taste-responsive and non-taste-responsive neurons of each cell type (**Figure 4B**; pyramidal cells: n = 65 taste-responsive cells, n = 376 non-taste-responsive cells; interneurons: n = 31 taste-responsive cells, n = 10 non-taste-responsive cells).

**Figure 2.**
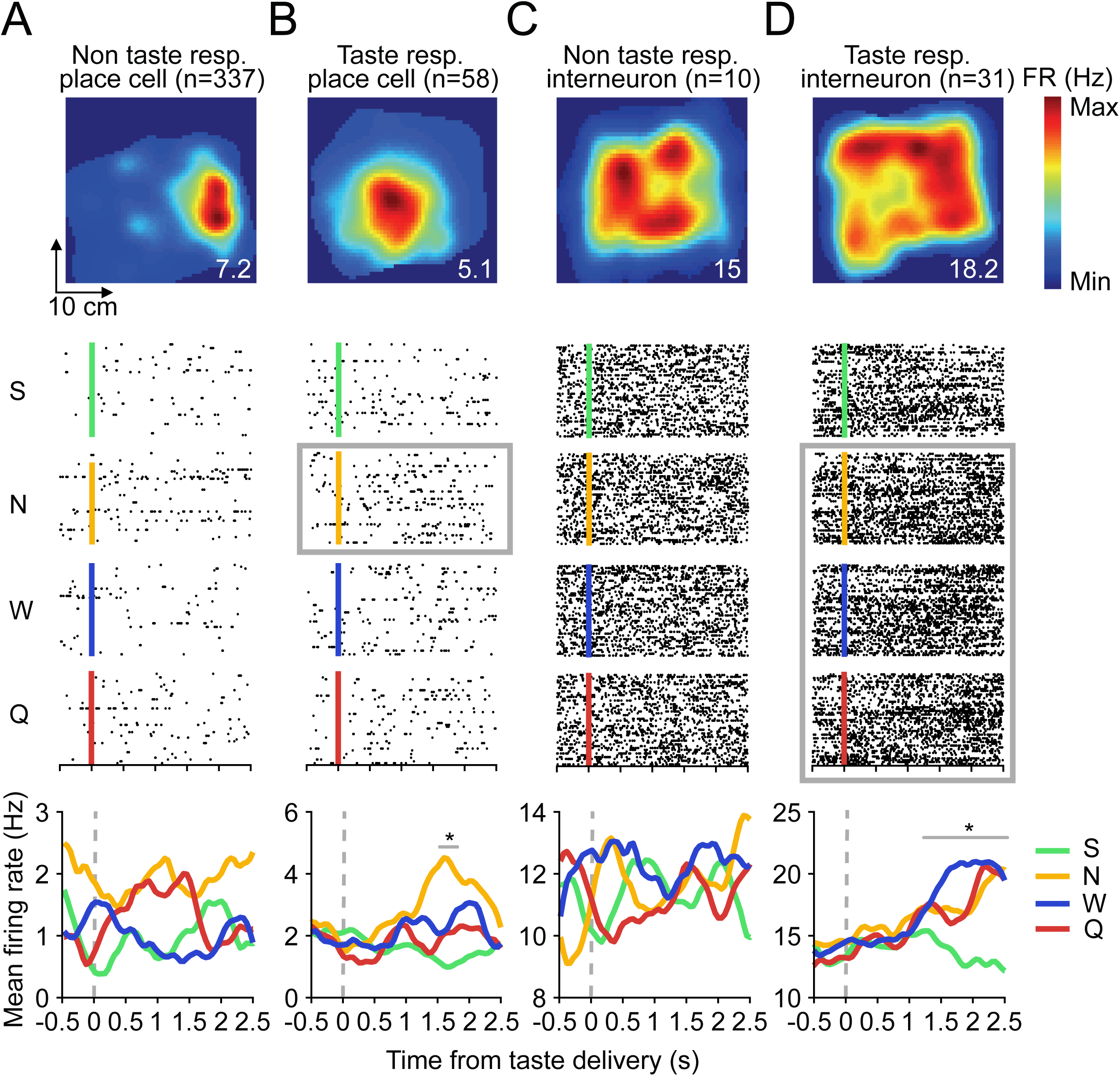
Subsets of hippocampal place cells and interneurons respond to tastes. ***A-D***, Top panels, Example spatial firing maps of two place cells (left) and interneurons (right), calculated with taste delivery periods (500 ms before to 2500 ms after taste delivery) omitted from the analysis. Numbers on the bottom right of each plot denote peak spatial firing rate (FR) in Hz. Middle panels, Raster plots of the cells in **A-D** responding to each of the four tastes, with trials aligned to the time of taste delivery (S, green line, 4 mM saccharin; N, yellow line, 100 mM sodium chloride; W, blue line, distilled water; Q, red line, 5 mM quinine hydrochloride) and black dots indicating when spikes occurred during each trial. Light gray boxes indicate the taste(s) for which there was a significant evoked response (*p < 0.05, t-tests on successive time windows). Bottom panels, Taste-evoked responses of each of the above neurons. Each colored trace represents the mean firing rate to one of the four tastes, smoothed with a 1D Gaussian filter (σ= 5 ms). Light gray lines indicate the periods of significant taste responsiveness (*p < 0.05, t-tests on successive time windows) for the place cell in **B** and interneuron in **D**.

Spatial specificity was determined by calculating the spatial information content, or amount of information that a single spike conveys about the animal’s location in bits/spike using the formula:

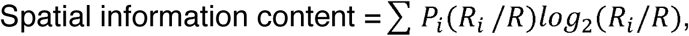

where *i* is the bin number, P_i_ is the probability of occupancy for bin *i*, R_i_ is the mean firing rate for bin *i*, and R is the overall mean firing rate of the cell (Skaggs et al., 1993).

Unpaired t-tests were used to determine whether the average spatial information content differed significantly between taste-responsive and non-taste-responsive neurons of each cell type (**Figure 4C**; pyramidal cells: n = 65 taste-responsive cells, n = 376 non-taste-responsive cells; interneurons: n = 31 taste-responsive cells, n = 10 non-taste-responsive cells).

#### In-field vs. out-of-field analysis

To analyze how place cells responded to tastes delivered inside or outside of their place fields (**Figure 5**), only pyramidal cells exhibiting place-specific activity (n = 395 cells, defined as neurons whose peak rate exceeded 1 Hz and spatial information content exceeded 0.2 bits/spike, similar to Moita et al., 2003) were considered. Only place cells that contained at least ten in-field and out-of-field deliveries of each taste were included in this in-field vs. out-of-field analysis (n = 26 taste-responsive cells, n = 153 non-taste-responsive cells). A one-way ANOVA was used to assess differences between the mean in-field and out-of-field taste response magnitude (calculated using *η* ^2^, see *Taste selectivity*) of taste-responsive and non-taste-responsive cells (**Figure 5C**). To ensure that comparable numbers of trials of each taste were delivered in-field, a one-way ANOVA was used to compare the mean number of in-field trials for each of the four tastes. An unpaired t-test was also used to compare the total number of in-field trials for taste-responsive and non-taste-responsive cells. To determine whether numbers of taste-responsive cells were underreported due to inter-trial variability, we repeated our initial analyses of taste responsiveness (see *Taste response properties*) using only in-field trials of place cells that met the in-field vs. out-of-field analysis criteria (n = 179 cells). A chi-square test was used to evaluate whether this approach resulted in comparable numbers of presence-, identity-and palatability-responsive cells.

#### Taste response properties

The pseudo-randomized taste delivery paradigm used to characterize hippocampal responses to tastes is described above (see *Passive taste administration paradigm*). Taste responses were characterized separately for each of the 482 isolated neurons, focusing on the 2500 ms of spiking activity following each taste delivery, a time period that includes previously identified taste-related responses, but precedes swallowing behaviors that remove tastes from the tongue and make neural responses difficult to interpret (Travers and Norgren, 1986; Katz et al., 2001). We analyzed a set of response properties ranging from general to specific, as have been identified in other parts of the taste system, including the GC (Katz et al., 2001; Sadacca et al., 2012), BLA (Fontanini et al., 2009; Piette et al., 2012), and LH (Li et al., 2013). Neurons were classified as “taste-responsive” (see **Table 1** for summary) if they exhibited responses to taste presence, identity and/or palatability, as described below. All other neurons were classified as “non-taste-responsive.” All statistical tests were performed in MATLAB (MathWorks, Natick, MA, RRID: SCR_001622) and evaluated at a level of α = 0.05 unless otherwise specified, with a Bonferroni correction applied for multiple comparisons.

First, non-specific responses to taste presence (**Figure 6B**, light gray lines), which are common across all four types of taste delivery and thought to originate from somatosensory responses detecting a taste on the tongue, were determined by assessing whether evoked responses differed significantly from the baseline firing rate in responses collated across all 200 taste delivery trials (Katz et al., 2001). The significance of the difference was first established using the main effect for time in a two-way, mixed-effect ANOVA (taste [saccharin, NaCl, water, quinine] x time [successive 500 ms bins of firing rate]).

Next, responses to taste identity (**Figure 6B**, dark gray lines), in which at least one taste can be discriminated from the others, were assessed by determining if the evoked responses to the four tastes (this time, collated across the 50 deliveries of each unique taste) differed from each other. We employed a similar strategy as the one used to evaluate taste responsiveness, except in this case, the main effect for taste was considered.

Finally, responses to taste palatability (**Figure 6D**), which reflected the relative hedonic value of tastes as assessed in the BAT (see **Figure 6C** and *Palatability/preference data*) were computed using a Spearman rank-order correlation (r_s_) between the evoked response and the palatability of the associated taste. Specifically, neurons whose evoked firing rates matched the ranking of taste preference (S > N > W > Q) in increasing or decreasing order had higher palatability index scores.

#### Population responses to tastes

For all taste-responsive cells (n = 96, see *Taste response properties* for classification criteria), the mean evoked firing rate across trials was determined (window size, 500 ms; step size, 50 ms; span, 0-2500 ms after taste delivery; Sadacca et al., 2012; Li et al., 2013) and Z-scored separately for each taste (**Figure 3A**). Responses to all four tastes were sorted by the timing of their peak firing to saccharin.

**Figure 3.**
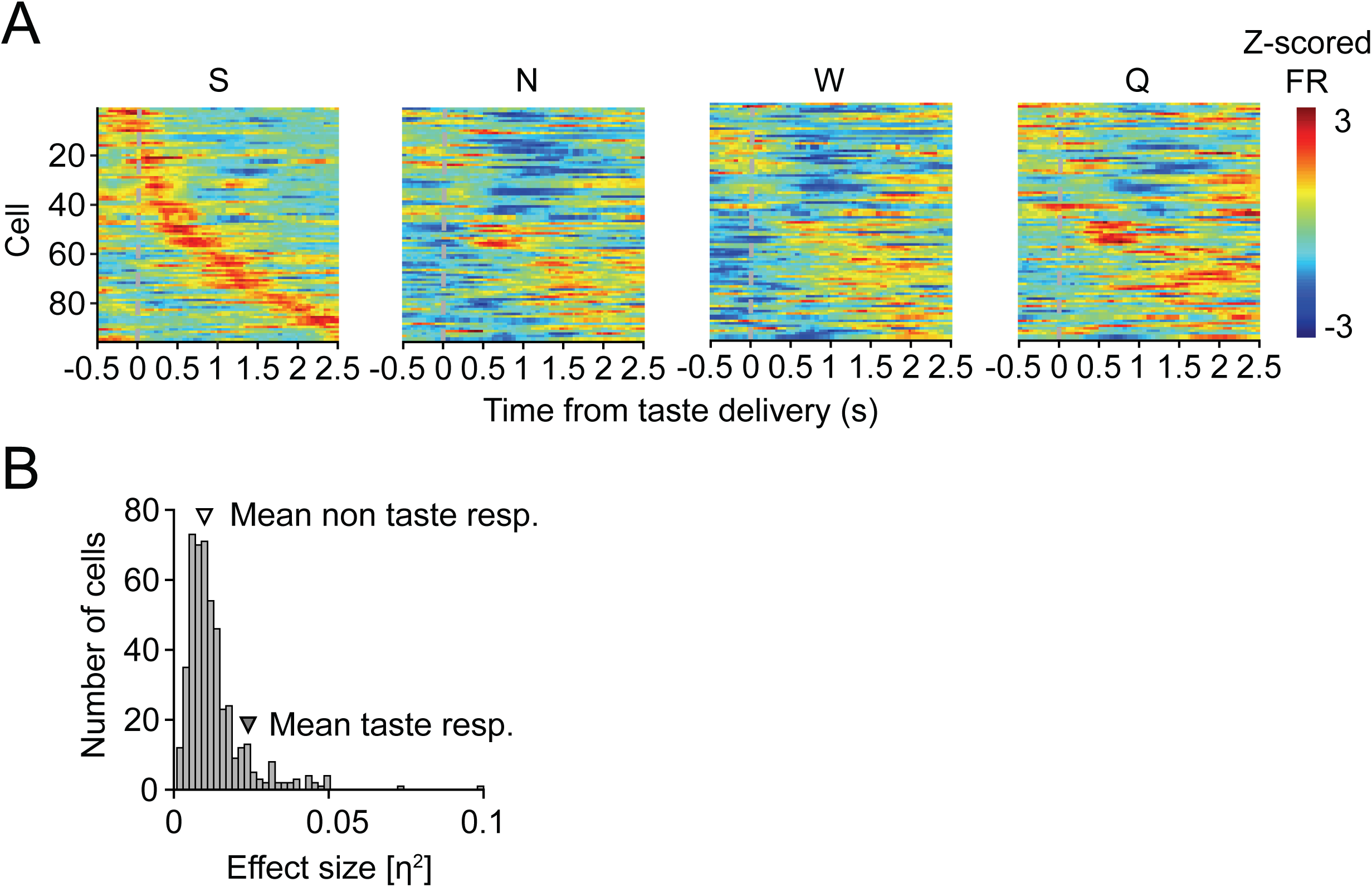
Diversity among the population of taste-responsive cells. ***A***, Z-scored firing rates of the 96 taste-responsive cells, ordered by the timing of their peak response to saccharin (S). Note the dissimilarity of responses when Z-scored firing rates to NaCl (N), water (W), or quinine (Q) are depicted using the same order, indicating that different cells respond preferentially and dynamically to each of the four administered tastes. ***B***, Histogram depicting the non-normal distribution of taste response magnitudes (quantified using effect size, *η* ^2^) within the entire population of neurons (n = 482 cells; chi-square goodness-of-fit test, X^2^ = 57.5, ***p = 3.3e-14), as well as the average ^2^ value for taste-responsive (n = 386 cells; mean *η* ^2^ = 0.024 ± 1.6e-04) and non-taste-responsive cells (n = 96 cells; mean *η* ^2^ = 0.010 ± 1.3e-05) within this distribution.

#### Taste selectivity

The magnitude of taste responsiveness for each cell was quantified using eta-squared (*η* ^2^), a standard measure of ANOVA effect sizes that describes the proportion of variance in a dependent variable explained by each factor:

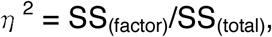

where SS is the sum of squares (Maier et al., 2015). In our analysis, we used the summed SS of the two main factors (time + taste) to calculate *η* ^2^. A chi-square goodness-of-fit test was used to determine if *η* ^2^ values followed a normal distribution (**Figure 3B**). The Pearson correlation (R) between spatial information content and *η* ^2^ was computed separately for place cells (n = 395 cells) and interneurons (n = 41 cells). As described above (see *In-field vs. out-of-field analysis*), a one-way ANOVA was used to assess differences in *η*^2^ for the in-field and out-of-field regions of taste-responsive (n = 26 cells) and non-taste-responsive (n = 153 cells) place cells that fit our analysis criteria (**Figure 5C**).

#### Taste response dynamics

To determine the timing of presence-, identity- and palatability-related responses in single neurons, Student’s t-tests were conducted on successive time windows of each neuron’s evoked response (window size, 500 ms; step size, 50 ms; span, 0-2500 ms after taste delivery; Sadacca et al., 2012; Li et al., 2013). To determine which taste(s) each of the identity- and palatability-responsive neurons (n = 40) preferentially responded to, the taste-evoked and baseline (−500 to 0 ms before taste delivery) firing rates were compared using successive t-tests. A chi-square goodness-of-fit test was used to determine whether cells responded to one taste, or to a particular number of tastes, more than the others.

To analyze taste-related dynamics on a population-wide level, we constructed a histogram showing what percentage of the total 482 recorded neurons exhibited responses to taste presence, identity and palatability at each time point following stimulus delivery (**Figure 7A**). To investigate the timing of different aspects of the taste experience present in hippocampal responses, we compared response onset times of presence- and identity-related firing (**Figure 7B**) as well as identity- and palatability-related firing (**Figure 7C**) in the subset of cells that exhibited both (n = 19 and 14 cells, respectively). Principal component analysis (PCA, see Briggman et al., 2005; Harvey et al., 2012) was conducted on the pooled subset of 36 identity-responsive cells to determine when discriminative firing emerged in the population response following taste delivery, with significance assessed at the α = 0.01 level comparing the neural data to 10,000 instances of firing-rate-shuffled controls (**Figure 7D**).

#### Speed and position controls

To ensure that hippocampal responses to tastes were not actually caused by overall differences in movement following taste delivery or in response to different tastes, we used a one-way ANOVA to compare the average speed and distance traveled during the pre-vs. post-taste period (2.5 seconds before or after taste delivery, segmented into 500 ms bins with a 50 ms step size) across all tastes (n = 1800 total trials across 9 sessions), as well as separately for each of the four tastes (n = 450 trials of each taste across 9 sessions). To confirm that tastes were delivered in different spatial locations in our paradigm, we used a one-way ANOVA to compare the mean number of taste trials in each spatial quadrant of the behavioral chamber.

## Results

### Hippocampal place cells and interneurons respond to tastes

We examined taste responses in a total of 482 CA1 neurons recorded across nine sessions in five rats (mean ± SEM: 53.6 ± 5.34 neurons/session) that received a battery of four standard tastes *via* IOC (**Figure 1A**). Tastes were delivered in random order and timing as rats explored the behavioral chamber, leading to a varied distribution of taste delivery locations, as exemplified in **Figure 1B** (mean number of trials across sessions, quadrant 1: 46.4 ± 6.57 trials, quadrant 2: 55.2 ± 14.58 trials, quadrant 3: 64.2 ± 12.16 trials, quadrant 4: 34.1 ± 5.34 trials; one-way ANOVA, p = 0.23). Histology confirmed that the majority of our tetrodes were located intermediately along the proximodistal axis of dorsal CA1 (**Figure 1C**; Henriksen et al., 2010). Isolated single neurons were classified as either putative pyramidal cells (91.5%, 441/482) or interneurons (8.5%, 41/482) on the basis of baseline firing rates and action potential shape (**Figure 1D**).

In total, 395 of the 441 pyramidal neurons were classified as place cells using standard analysis of the spatial specificity of firing rate responses (see Materials and Methods, Moita et al., 2003). The spatial firing maps of four representative place cells and interneurons (all of which were computed with taste delivery periods omitted from the analysis) are shown in the top row of **Figure 2**. As expected (O’Keefe and Dostrovsky, 1971; O’Keefe and Nadel, 1978), only the pyramidal cells had place fields (**Figure 2A, 2B**)—interneurons (**Figure 2C, 2D**) typically exhibited high spontaneous firing rates regardless of the rat’s position.

A cell was considered “taste-responsive” if significant firing rate modulations were evoked by taste presence, identity, and/or palatability (see **Figure 6** for more details). In total, 96/482 (19.9%) cells were classified as taste-responsive, which is similar to the proportion reported in the only previous study to assess taste responses in individual hippocampal neurons (Ho et al., 2011). We found taste-responsive and unresponsive units on tetrodes across the proximodistal axis of dorsal CA1 (n = 50/60 tetrodes with taste-responsive units). **Table 1** shows the distribution of taste-responsive cells across animals.

The raster plots and peri-stimulus time histograms (PSTHs) for an example taste-responsive place cell and interneuron are depicted in **Figure 2B** and **2D**. The place cell in **Figure 2B** responded to NaCl from 1550-1750 ms following taste delivery, while the interneuron in **Figure 2D** responded to NaCl, water and quinine from 1200-2500 ms following taste delivery (light gray lines). In contrast, the raster plots and PSTHs for non-taste-responsive cells (**Figure 2A, 2C**) show no differences in evoked activity from baseline or between tastes.

The Z-scored firing rates (**Figure 3A**) reveal the diversity of taste responses within the population of taste-responsive cells (n = 96 cells). When Z-scored firing rates are ordered by the timing of each cell’s peak excitatory response to saccharin (S, **Figure 3A**), these responses tile the entirety of the taste delivery period. The breakdown of this sequence for NaCl (N), water (W), and quinine (Q) when the same cell order is conserved illustrates that taste-responsive cells respond preferentially and dynamically to different tastes. **Figure 3B** reveals that the distribution of taste response magnitudes as quantified by eta-squared (*η* ^2^, a standard measure of ANOVA effect size; see Maier et al., 2015) does not follow a normal distribution (number of histogram bins = 50, bin width = 2e-04; chi-square goodness-of-fit test, X ^2^ = 57.5, p = 3.3e-14). Instead, the majority of neurons (n = 386 cells) were non-taste-responsive and clustered around low *η* ^2^ values, while a smaller proportion of neurons (n = 96 cells) were taste-responsive and had higher *η* ^2^ values (mean *η* ^2^, taste-responsive cells: 0.024 ± 1.6e-04, non-taste-responsive cells: 0.010 ± 1.3e-05).

Since hippocampal activity is affected by animals’ location and movement, one possible explanation of these results is that different tastes have different impacts on animals’ motor behavior, and that therefore any perceived “taste”-evoked responses can simply result from changes in the animal’s speed or position (O’Keefe, 1999; Shan et al., 2016). To control for this possibility, we assessed differences in rats’ pre- and post-taste speed and position, both overall and between each of the four tastes. We found no differences in the average speed (before taste delivery: 1.07 ± 0.031 cm/s, after taste delivery: 1.12 ± 0.029 cm/s; one-way ANOVA, p = 0.29) or distance traveled (before taste delivery: 1.57 ± 0.050 cm, after taste delivery: 1.59 ± 0.045 cm; one-way ANOVA, p = 0.74) in the 2.5 seconds preceding and following taste deliveries; the same was true when trials were split up by taste identity (post-pre taste delivery speed, saccharin: −0.081 ± 0.063 cm/s, NaCl: 0.037 ± 0.061 cm/s, quinine: 0.11 ± 0.061 cm/s, water: 0.12 ± 0.070 cm/s; one-way ANOVA, p = 0.10; post-pre taste delivery distance: saccharin: −0.0026 ± 0.11 cm, NaCl: −0.029 ± 0.11 cm, quinine: 0.093 ± 0.093 cm, water: 0.027 ± 0.11 cm; one-way ANOVA, p = 0.86). Therefore, it is unlikely that hippocampal responses to tastes were caused by changes in animals’ position or locomotion; rather, they reflected sensory responses to some aspect of the taste experience itself.

### Taste responses are gated by the spatial firing properties of hippocampal neurons

We found that 14.7% of place cells (n = 58/395 cells) had significant responses to tastes; a far-higher percentage of interneurons (75.6%; n = 31/41 cells) were taste-responsive (**Figure 2**). The significance of this larger likelihood of taste-responsiveness amongst spatially diffuse interneurons than in spatially-specific place cells (chi-square test, X^2^ = 84.87, p = 3.19e-20) suggests that taste responsiveness depends on the spatial firing properties of hippocampal neurons. To further investigate the relationship between place- and taste-specific firing, we compared the place field size (Brun et al., 2002), spatial information content (Skaggs et al., 1993), and taste response magnitudes (Maier et al., 2015) of taste-responsive and non-taste-responsive hippocampal neurons.

**Figure 4A** depicts the firing fields of 12 example non-taste-responsive and taste-responsive place cells and interneurons, all of which were computed with taste delivery periods (extending from 500 ms before to 2500 ms after taste delivery) omitted from the analysis. As expected, place cells had a smaller firing field size (mean field size, place cells: 525.1 ± 18.1 cm^2^, interneurons: 988.7 ± 46.4 cm^2^; unpaired t-test, p = 1.4e-14) and higher spatial information content than interneurons (mean spatial information content, place cells: 1.3 ± 0.034 bits/spike, interneurons: 0.12 ± 0.034 bits/spike; unpaired t-test, p = 2.6e-25; higher values = smaller, more concentrated regions of enhanced firing). Therefore, taste responses (which were found predominately in interneurons) were associated with less spatially selective firing.

**Figure 4.**
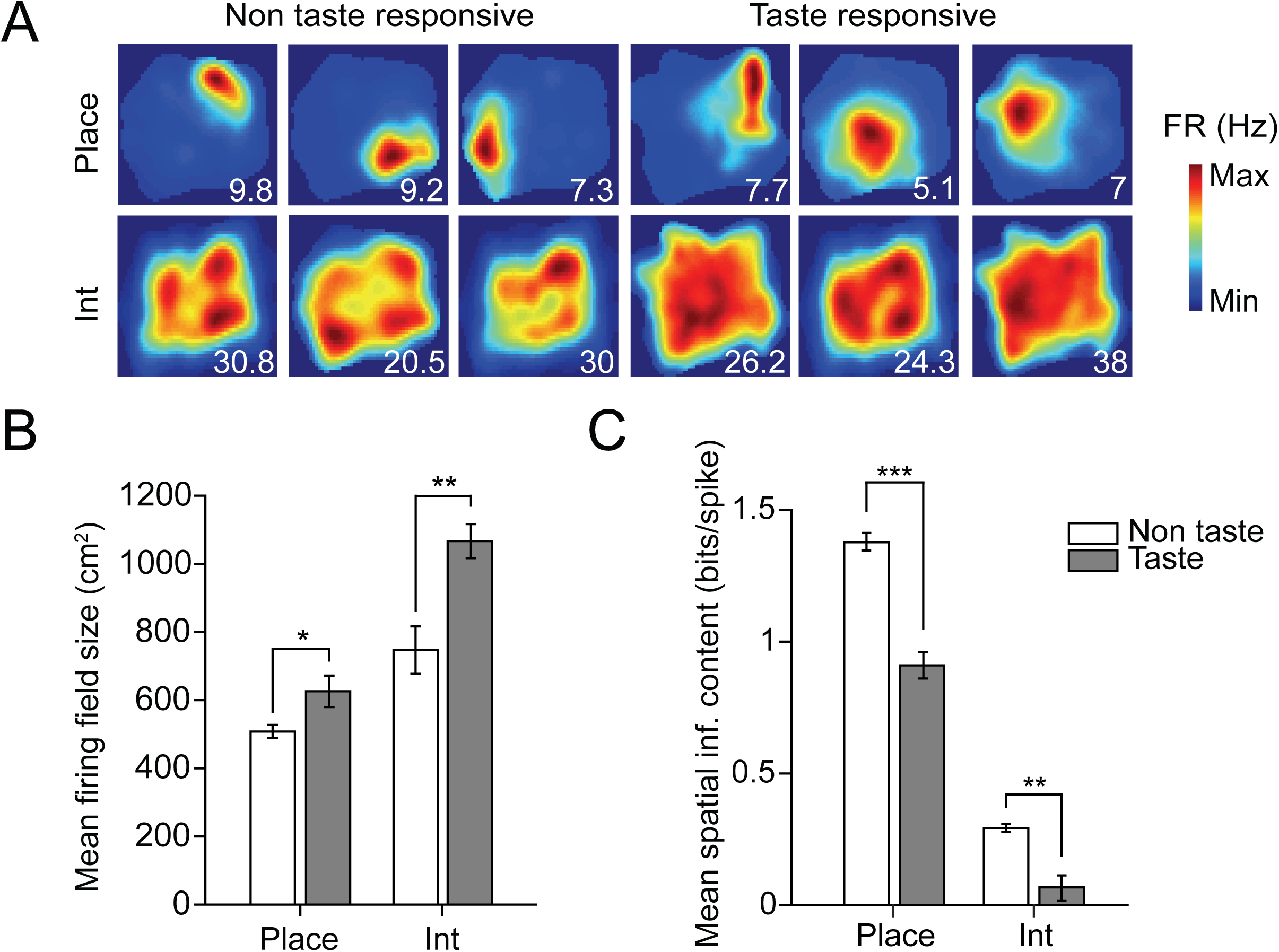
Taste-responsive hippocampal neurons exhibit weaker spatial selectivity than non-taste-responsive hippocampal neurons. ***A***, Example spatial firing maps of twelve non-taste-responsive (left) and taste-responsive (right) place cells (Place, top row) and interneurons (Int, bottom row). Note that taste-responsive cells tend to exhibit more diffuse spatial firing. All firing maps were computed with taste delivery periods (500 ms before to 2500 ms after taste delivery) omitted from the analysis. Numbers on the bottom right of each plot denote peak spatial firing rate (FR) in Hz. ***B***, Mean firing field size for non-taste-response (white bars) and taste-responsive (gray bars) place cells and interneurons. For each cell type, taste-responsive neurons had larger firing fields than non-taste-responsive neurons (place cells: n = 337 non-taste-responsive cells, n = 58 taste-responsive cells; unpaired t-test, *p = 0.021; interneurons: n = 10 non-taste-responsive cells, n = 31 taste-responsive cells; unpaired t-test, **p = 0.002). ***C***, Mean spatial information content for non-taste-responsive (white bars) and taste-responsive (gray bars) place cells and interneurons. Within each cell type, taste-responsive neurons had a lower spatial information content than non-taste-responsive neurons (place cells, unpaired t-test, ***p = 1.03e-06; interneurons, unpaired t-test, **p = 0.0027).

This same pattern was found to hold even within each cell type, however: cells with stronger taste-evoked responses tended to have larger place fields (**Figure 4B**) and exhibit weaker spatial responses (**Figure 4C**) in analyses restricted to place cells (mean firing field size, taste-responsive place cells: 626.0 ± 46.1 cm^2^, non-taste-responsive place cells: 507.7 ± 19.6 cm^2^; unpaired t-test, p = 0.021; mean spatial information content, taste-responsive place cells: 0.91 ± 0.050 bits/spike, non-taste-responsive place cells: 1.37 ± 0.038 bits/spike; unpaired t-test, p = 1.03 e-06) as well as interneurons (mean firing field size, taste-responsive interneurons: 1066.7 ± 49.9 cm^2^, non-taste-responsive interneurons: 746.8 ± 69.7 cm^2^; unpaired t-test, p = 0.002; mean spatial information content, taste-responsive interneurons: 0.065 ± 0.040 bits/spike, non-taste-responsive interneurons: 0.29 ± 0.015 bits/spike; unpaired t-test, p = 0.0027). There was a negative correlation between the spatial information content and magnitude of taste responsiveness (*η* ^2^) within each cell type (place cells: Pearson correlation, R = −0.18, p = 3.27e-04; interneurons: Pearson correlation, R = −0.58, p = 5.95e-05), confirming that hippocampal neurons that respond strongly to taste delivery tend to have more diffuse firing in space.

The above analysis implies that while place cells tended to exhibit fewer and lower-magnitude taste responses than interneurons, a subset of place cells exhibited taste-specific firing (n = 58/395 cells; example in **Figure 2B**). Thus it is important to ask how place and taste responses interact when an animal receives familiar tastes in a specific spatial context: can place cells acquire sensory responses regardless of location, or are responses to tastes gated by spatial firing, as suggested for other sensory modalities (Moita et al., 2003; Shan et al., 2016)? To investigate this question, we compared the specificity of taste responses inside and outside each place cell’s firing field.

Only place cells (n = 58 taste-responsive cells, n = 337 non-taste-responsive cells) were considered for in-field vs. out-of-field analysis. We defined a cell’s place field as the largest number of adjacent spatial bins which had firing rates >= 20% of the peak firing rate (Brun et al., 2002). To ensure sufficient sampling of taste responses for statistical comparisons, only cells that contained at least ten in-field and out-of-field trials of each taste stimulus were included in our analysis (n = 26 taste-responsive cells, n = 153 non-taste-responsive cells). Taste response magnitude (*η* ^2^) was then determined separately for trials taking place within and outside each cell’s place field.

The top row of **Figure 5** shows the spatial firing maps of representative taste-responsive (**Figure 5A**) and non-taste-responsive (**Figure 5B**) place cells, with in-field and out-of-field trials indicated in pink. The distribution of taste trials in these cells was representative of the entire population analyzed, with comparable numbers of taste trials delivered in-field (number of in-field trials, taste-responsive cells, saccharin (S): 25.1 ± 1.5 trials, NaCl (N): 25.5 ± 1.5 trials, water (W): 24.4 ± 1.4 trials, quinine (Q): 24.8 ± 1.5 trials; one-way ANOVA, p = 0.96; number of in-field trials, non-taste-responsive cells, saccharin: 23.4 ± 0.7 trials, NaCl: 24.1 ± 0.7 trials, water: 23.3 ± 0.7 trials, quinine: 23.8 ± 0.7 trials; one-way ANOVA, p = 0.89). The total number of in-field trials was also comparable across groups (number of in-field trials, taste-responsive cells: 99.7 ± 5.7 trials, non-taste-responsive cells: 94.6 ± 2.7 trials; unpaired t-test, p = 0.46). Repeating our initial analyses of taste responsiveness using only the in-field trials of units that met our analysis criteria (n = 179 cells) resulted in comparable numbers of taste-responsive cells (n = 17 cells using in-field trials, n = 26 cells using all trials; chi-square test, X^2^ = 1.69, p = 0.19), indicating that the total number of taste-responsive neurons was not underestimated due to inter-trial variability, and cells previously classified as non-taste-responsive did not typically contain any regions of taste-responsiveness, either in- or out-of-field (see example in **Figure 5B**).

**Figure 5.**
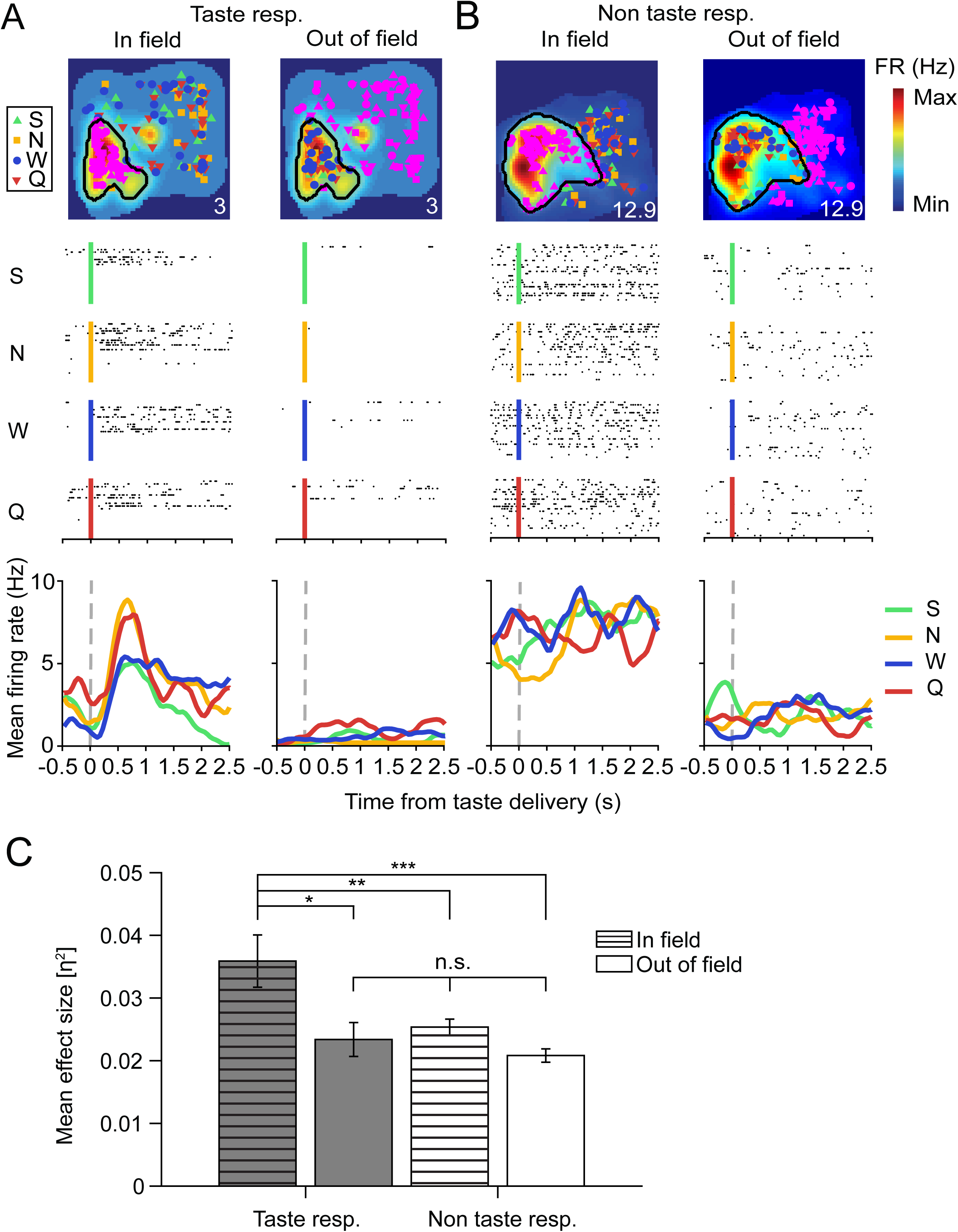
Place cells respond to tastes delivered within their place field. ***A-B***, Example in-field and out-of-field responses for a taste-responsive (**A**) and non-taste-responsive (**B**) place cell. Top panels show the spatial firing maps for each cell, with place field boundaries (defined as largest cluster of neighboring spatial bins which had firing rates >= 20% of the peak rate) indicated by black lines. The colored symbols represent locations of individual taste deliveries (green: S, 4 mM saccharin; yellow: N, 100 mM sodium chloride; blue: W, distilled water; red: Q, 5 mM quinine hydrochloride), with either in-field (left) or out-of-field (right) trials highlighted in pink. All firing maps were computed with taste delivery periods (500 ms before to 2500 ms after taste delivery) omitted from the analysis. Numbers on the bottom right of each plot denote peak spatial firing rate (FR) in Hz. Middle panels depict raster plots of each cell’s in-field (left) and out-of-field (right) responses to each of the four tastes. Black dots indicate spike times during each individual trial, aligned to the time of taste delivery (S, green line, 4 mM saccharin; N, yellow line, 100 mM sodium chloride; W, blue line, distilled water; Q, red line, 5 mM quinine hydrochloride). Bottom panels show the evoked responses to taste deliveries taking place in-field and out-of-field for each cell. Each colored trace represents the mean firing rate to one of the four tastes, smoothed with a 1D Gaussian filter (σ= 5 ms). Note that taste responses are only found within the taste-responsive cell’s place field (left panel of **A**). ***C***, Mean magnitude of taste responsiveness (eta-squared, or *η* ^2^) for the in-field and out-of-field regions of taste-responsive and non-taste-responsive place cells that fit our criteria (>= 10 trials in- and out-of-field; n = 26 taste-responsive cells, n = 153 non-taste-responsive cells). The mean *η* ^2^value for the in-field region of taste-responsive place cells (striped gray bar) was higher than that of the in-field region of non-taste-responsive cells (striped white bar), the out-of-field region of taste-responsive cells (gray bar), or the out-of-field region of non-taste-responsive (white bar) cells (1-way ANOVA, ***p = 3e-05), indicating that place cells only respond to tastes delivered within their place field.

For the taste-responsive cell (**Figure 5A**), virtually all responses occurred within the cell’s place field (left panel), with very little taste-evoked firing out of field (right panel). On the other hand, no taste-evoked responses were observed in- or out-of-field for the non-taste-responsive cell (**Figure 5B**). This trend was representative of the entire population of place cells (**Figure 5C**): the in-field region of taste-responsive cells had a higher mean eta-squared value than the out-of-field region, or either region of non-taste-responsive cells (mean eta-squared values, taste-responsive cells, in-field: 0.036 ± 0.0042; taste-responsive cells, out-of-field: 0.023 ± 0.0027; non-taste-responsive cells, in-field: 0.025 ± 0.0013; non-taste-responsive cells, out-of-field: 0.0208 ± 0.0011; 1-way ANOVA, p = 3e-05). Together, these results indicate that hippocampal taste responses are gated by the spatial firing properties of place cells—a finding that is consistent with previous studies investigating tone-evoked sensory responses during auditory fear conditioning (Moita et al., 2003; Shan et al., 2016).

### Hippocampal taste-specific responses reflect taste palatability, at a relatively long delay

Previous work has shown that taste-specific firing in GC neurons evolves through three stages: following an initial, nonspecific response to taste presence, a discriminative response conveys information about taste identity starting at approximately 200 ms after stimulus administration; after approximately 500 ms, responses then change to reflect palatability, specifically anticipating an animal’s decision to consume or expel a particular taste between 600 and 1600 ms after taste delivery (Katz et al., 2001; Piette et al., 2012; Sadacca et al., 2012; Maier and Katz, 2013; Li et al., 2016; Sadacca et al., 2016). Brainstem taste responses in the parabrachial nucleus (PbN) are similarly organized (Baez-Santiago et al., 2016). Other nodes of the taste CNS, however, such as the BLA and LH, appear instead to respond primarily and immediately to the hedonic value of tastes, regardless of identity (Nishijo et al., 1998; Fontanini et al., 2009; Li et al., 2013). To determine which components are present in hippocampal taste responses (and when), we performed analyses similar to those brought to bear on firing in these other structures.

Many hippocampal neurons responded nonspecifically to taste presence, providing information that could allow for the detection of tastants on the tongue (**Figure 6A**). In a subset of these cells, responses were more discriminative, providing information about taste identity and/or palatability. Two such neurons are shown in **Figure 6B**—one of which (place cell, left) rapidly developed a response primarily to quinine, and one of which (interneuron, right) produced a longer-latency response that differentiated each of the four tastes, and which notably involved a sudden change of firing rate to saccharin. As illustrated previously (**Figure 3A**), we observed a relatively uniform distribution among taste-discriminative cells (n = 40) of which taste(s) cells responded to (number of cells with significant evoked responses to saccharin: 20 cells, NaCl: 29 cells, water: 26 cells, quinine: 25 cells; chi-square goodness-of-fit test, X^2^ = 1.7, p = 0.64). Similar to the interneuron shown in **Figure 6B**, most taste-discriminative cells responded to more than one taste (number of cells responding to one taste: 11 cells, two tastes: 4 cells, three tastes: 19 cells, four tastes: 6 cells; chi-square goodness-of-fit test, X^2^ = 13.4, p = 0.0038).

**Figure 6.**
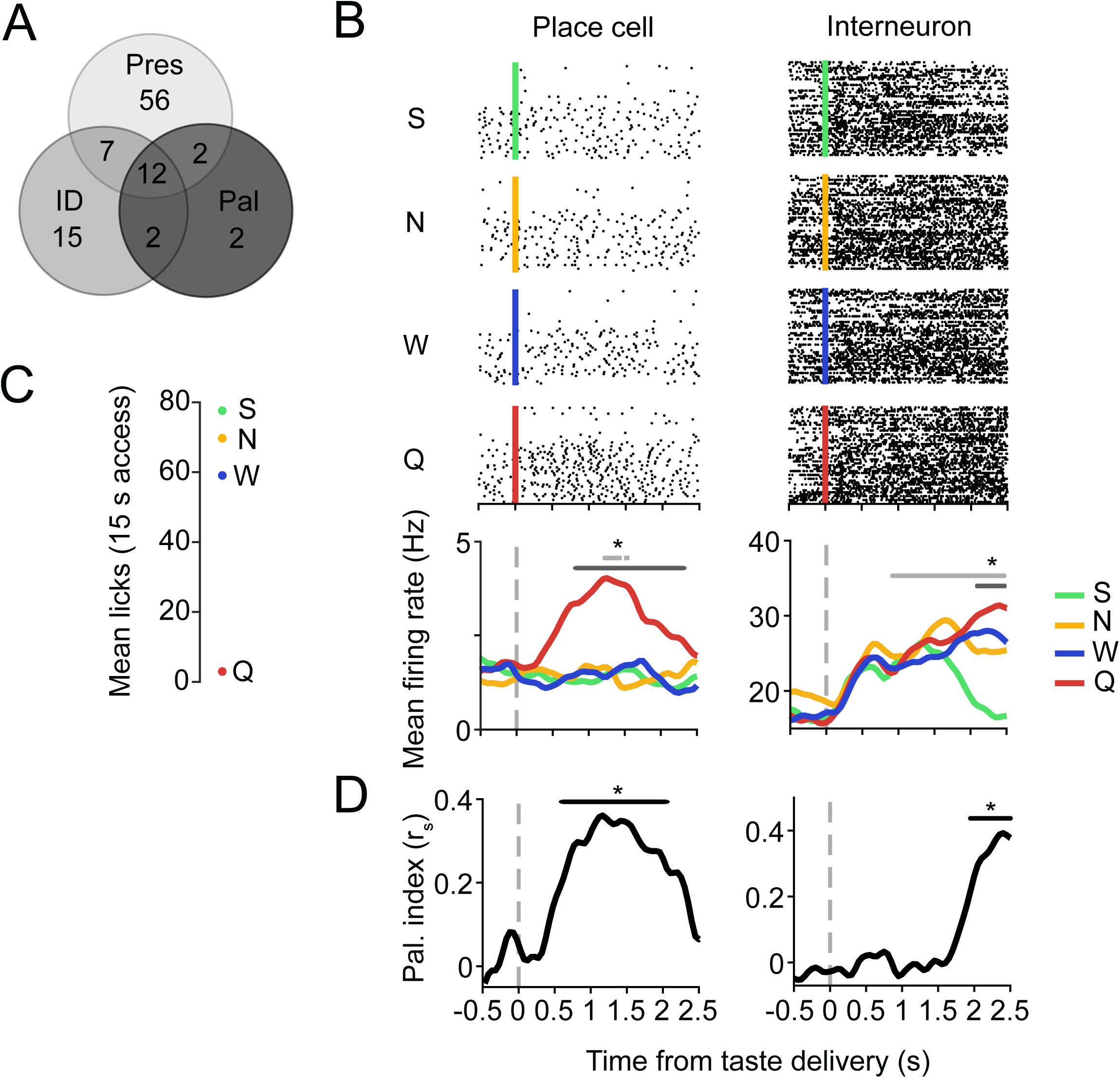
Example hippocampal responses to different elements of the taste experience. ***A***, Summary of the number of taste-responsive cells (n = 96/482 cells) that responded to taste presence (Pres, n = 77 cells), identity (ID, n = 36 cells), and/or palatability (Pal, n = 18 cells). ***B***, Example raster plots and PSTHs from a taste-responsive place cell (left) and interneuron (right). Black dots in the raster plots (top four panels) represent spike times during each trial, aligned to taste delivery time (S, green line, 4 mM saccharin; N, yellow line, 100 mM sodium chloride; W, blue line, distilled water; Q, red line, 5 mM quinine hydrochloride). Each colored trace in the PSTHs (bottom panels) represents the mean firing rate to one of the four tastes, smoothed with a 1D Gaussian filter (σ= 5 ms). Light gray lines indicate periods of significant responses to taste presence, while dark gray lines indicate periods of significant responses to taste identity (*p < 0.05, t-tests on successive time windows). ***C***, Relative palatability of the four taste stimuli as determined by a brief-access task (BAT). Palatability rank is determined by the average number of licks per 15 s of exposure to the given taste. ***D***, Rank-order correlation (r_s_) between the taste-evoked firing rates and palatability rank (S > N > W > Q) for the place cell (left) and interneuron (right) depicted in **B**. Black lines indicate periods of significant palatability-relatedness (*p < 0.05, t-tests on successive time windows). Note the similarity between the timing of palatability- and identity-related responses (dark gray lines in **B**).

Closer examination revealed that the patterning of both responses in **Figure 6B** reflected taste palatability across the entirety of the periods of taste-specific firing (**Figure 6D**). Responses to taste palatability were assessed, as is typical in studies of taste temporal coding (Li et al., 2013; Baez-Santiago et al., 2016; Sadacca et al., 2016), in terms of the correlation between neuronal firing rates and the order of taste preference, which was assayed in a brief-access task (Li et al., 2013; Monk et al., 2014; Sadacca et al., 2016) run on a separate cohort of experimental rats (**Figure 6C**). The observed order of taste preference (S > N > W > Q) shown in **Figure 6C** is consistent with that observed across a broad range of stimulus delivery methods and assessment techniques (Travers and Norgren, 1986; Breslin et al., 1992; Clarke and Ossenkopp, 1998; Fontanini and Katz, 2006; Sadacca et al., 2016).

**Figure 6D** reveals that palatability correlations for the example neurons shown in **Figure 6B** developed as the taste-specific responses themselves developed: the place cell’s responses (**Figure 6B**, left) were significantly correlated with taste palatability between 600 and 2100 ms (**Figure 6D**, left), while the interneuron’s responses (**Figure 6B**, right) were palatability-related between 2000 and 2500 ms (**Figure 6D**, right; compare these periods with the dark gray line in **Figure 6B**, which marks the period of significantly taste-specific firing).

Again, the examples shown in **Figure 6** suggest that, like responses observed in limbic structures (i.e., BLA and LH; Fontanini et al., 2009; Li et al., 2013) but unlike those in the main taste axis (i.e., GC and PbN; Sadacca et al., 2012; Baez-Santiago et al., 2016), hippocampal taste responses do not go through a period of “pure” taste specificity prior to becoming palatability-related. These appearances were borne out in an analysis of the entire neural dataset. **Figure 7A** shows, similar to what has been observed in all other parts of the taste system (Katz et al., 2001; Sadacca et al., 2012; Baez-Santiago et al., 2016; Fontanini et al; 2009; Li et al., 2013), that totally non-specific responses to taste presence emerged first in hippocampus, followed by responses to taste identity and palatability. However, both taste specificity and palatability-relatedness appeared in hippocampal taste responses at similarly long latencies (average onset, presence: 1032.5 ± 55.73 ms; identity: 1443.1 ± 108.07 ms; palatability: 1797.2 ± 118.51 ms).

**Figure 7.**
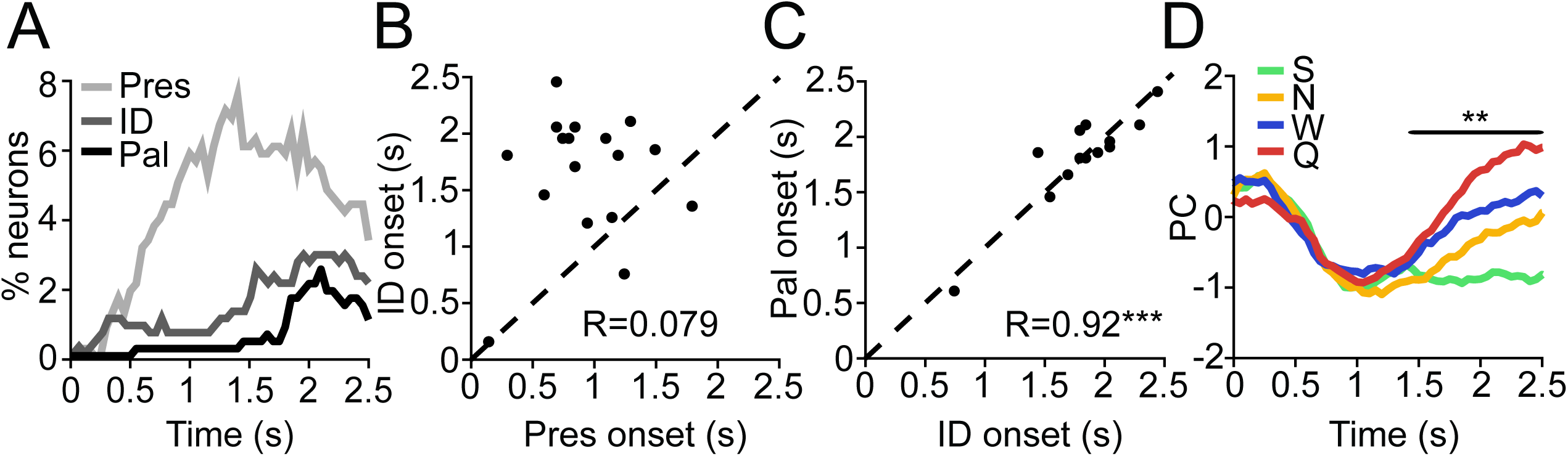
Hippocampal neurons discriminate between tastes based on palatability. ***A***, Histogram showing the percentage of hippocampal neurons that responded significantly to taste presence (Pres, light gray line), identity (ID, dark gray line) or palatability (Pal, black line) at each time point relative to taste delivery. Nonspecific responses to taste presence emerged before responses to taste identity or palatability, which occurred at similarly long latencies. ***B***, For the 19 cells that responded significantly to taste presence and identity, the onset of the identity-related response (y-axis) is plotted against the onset of the presence-related response (x-axis). Similar to what is seen in **A**, single-neuron responses to taste presence typically preceded responses to taste identity, as evidenced by the cloud of points above the unity line (black dashed line). ***C***, For the 14 cells that responded significantly to taste identity and palatability, the onset of the palatability-related response (y-axis) is plotted against the onset of the identity-related response (x-axis). Responses to identity and palatability tended to emerge simultaneously in single units, as evidenced by tight clustering around the unity line (regression slope = 0.92, ***p = 8.68e-06). ***D***, PCA of identity-responsive cells (n = 36 cells). Each colored line depicts the PC of pooled responses to each of the four tastants (green: S, 4 mM saccharin; yellow: N, 100 mM sodium chloride; blue: W, distilled water; red: Q, 5 mM quinine hydrochloride) over time. Palatability-relatedness emerges at the same time as taste selectivity, as shown by significant encoding of palatability rank (S > N > W > Q, in reverse order here; **p < 0.01, comparison to firing-rate-shuffled controls) by the PCs starting at ~1.4 s after stimulus delivery (black line).

Direct within-neuron comparisons strongly supported the group analysis. Presence-related responses reliably arose before identity-related responses in cells that responded to both properties (n = 19 cells, paired t-test, p = 7.6e-05), as also evidenced by the cloud of points above the unity line (**Figure 7B**; regression slope: 0.079, p = 0.79). A plot of the onset latency of identity- and palatability-related responses in cells where both properties were present, meanwhile, revealed tight clustering around the unity line (**Figure 7C**; n = 14 cells; regression slope: 0.92, p = 8.7e-06) with no significant differences between onset times (paired t-test, p = 0.83), suggesting that these properties arose simultaneously in single-unit responses.

Finally, we performed PCA by pooling responses of identity-responsive cells (n = 36 cells) to visualize the onset of taste-specific firing in the population as a whole. This analysis suggests that tastes are discriminated based on palatability, as shown by significant encoding of palatability rank (here, in reverse order as shown in **Figure 6C**) by the principal components starting at ~1.4 s after stimulus delivery (**Figure 7D**). It is important to note that this analysis is not representative of every cell: we identified a small but significant contingent of exclusively identity-coding neurons (n = 15/96 taste-responsive cells; see **Figure 6A**, and **Figure 2B** for example); it’s worth noting, however, that even many of the neurons that responded strongly and specifically to single tastes did so to either the most rewarding or aversive tastants in our battery (see **Figure 6B** and **D** for examples).

We therefore conclude that hippocampal “taste codes” do not contain the purely identity-related component found in gustatory brainstem and cortex; rather, taste selectivity emerges at the same time as palatability-relatedness, and as a whole, the taste-reactive hippocampal population delivers information on palatability without a prior epoch of less structured taste-specificity. In this regard, hippocampal responses are similar to those observed in other non-cortical parts of the taste system, such as the BLA (Fontanini et al., 2009) and LH (Li et al., 2013); notably, however, palatability coding appears in hippocampus much later than it appears in these other limbic structures—a difference that likely has strong implications for the potential roles of the hippocampus in taste (see Discussion below).

## Discussion

Our findings suggest that place and taste responses can co-exist within the same hippocampal neurons, and that these response modalities influence one another. Taste-responsive cells tended to have less spatially specific firing fields (**Figures 2**, **4**). On the other hand, place cells that responded to tastes did so in a spatially specific manner (**Figure 5**), with responses only occurring within that cell’s place field. Hippocampal neurons discriminated between tastes at relatively long latencies and predominantly on the basis of palatability (**Figures 6**, **7**), confirming that these responses can encode sensory parameters, and do not simply reflect changes in animals’ movement or attentive state, as has been suggested (O’Keefe, 1999; Shan et al., 2016). These results stand alongside similar studies of audition (Moita et al., 2003) and olfaction (Wood et al., 1999), and further show that spatially tuned hippocampal neurons can discriminate between behaviorally relevant sensory stimuli. Our observations add to an expanding view of the hippocampal cognitive map as a representation that encompasses both spatial and non-spatial aspects of an animal’s environment (Shapiro et al., 1997; Eichenbaum et al., 1999; Wood et al., 1999; Moita et al., 2003; Kraus et al., 2013; Aronov et al., 2017).

In total, about 20% of recorded hippocampal cells in our study were classified as taste-responsive (**Figure 2**), which is comparable to the proportion reported in the only previous study to assess taste responses in individual hippocampal neurons (Ho et al., 2011). Our usage of IOCs for enhanced stimulus control allowed us to build upon Ho and colleagues’ findings, uncovering new understanding of the structure, content, and dynamics of hippocampal taste responses. Similar to what is seen elsewhere in the taste system, including the GC (Katz et al., 2001), BLA (Fontanini et al., 2009), LH (Li et al., 2013), and PbN (Baez-Santiago et al., 2016), we observed larger-than-chance numbers of CA1 neurons that responded discriminatively to taste stimuli (**Figure 6A**), with response specificity evolving over time (**Figure 7**). This taste-distinctiveness, along with the history of taste research that has been conducted in rodents using IOCs, further underscores that hippocampal taste responses are truly gustatory, and do not arise from stress or fear responses caused by the delivery of solutions via IOC (Travers and Norgren, 1986; Katz et al., 2001; Sadacca et al., 2016).

Unlike what has been observed in GC (Katz et al., 2001; Sadacca et al., 2016) and PbN (Baez-Santiago et al., 2016), we found little evidence of pure identity coding in the hippocampus in this passive administration paradigm. Instead, hippocampal neurons distinguish between tastes based on palatability (**Figure 7**), similar to other taste system limbic structures such as the BLA (Fontanini et al., 2009) and LH (Li et al., 2013). However, palatability-related hippocampal coding emerged much later than in BLA or LH, and likely after the time (although more direct measurements must be taken to ascertain this) that animals make decisions about palatability-related orofacial behaviors (Li et al., 2016; Sadacca et al., 2016). These results support the idea that the hippocampus does not contribute to an animal’s decision to consume or expel a given taste. Rather, it responds to the hedonic value of tastes consumed within a particular context, forming associations between place and reward that may be relayed downstream via hippocampal projections to the ventral tegmental area and other neural reward centers (Lisman et al., 2011). This could serve as a means of relating place to taste in terms of its inherent reward value, allowing animals to use past experience to locate food sources.

While spatial learning is indisputably considered to be a hippocampal-dependent process (Morris, 1984; Moser et al., 2008), the role of the hippocampus in non-spatial taste learning is less clear-cut. Some forms of taste learning, such as social transmission of food preferences (Bunsey and Eichenbaum, 1995; Countryman et al., 2005) are considered to be hippocampal-dependent; other paradigms (e.g., conditioned taste aversion) undergo variable effects after hippocampal lesions (Yamamoto et al., 1995; Stone et al., 2005; Chinnakkaruppan et al., 2014). Future studies in which individual neurons are recorded during taste learning, which have been informative when focused on other nodes of the taste system (Grossman et al., 2008; Lavi et al., 2018), may help to decipher how the hippocampus encodes tastes and contexts to guide future food choices.

Our study provides the first direct evidence that hippocampal taste responses are almost entirely gated by the neurons’ spatial firing properties. More specifically, taste responses are more prevalent in interneurons than place cells (**Figure 2**), associated with broader spatial responsiveness regardless of cell type (**Figure 4**), and limited to cells’ spatial firing fields (**Figure 5**). Our finding that place cells only respond to tastes delivered within their place field is consistent with previous studies (Moita et al., 2003; Shan et al., 2016), indicating that taste responses can best be understood as a rate code overlaid on existing representations of space. Since place fields can be modulated by food reward (Dupret et al., 2010; Allen et al., 2012), it seems likely that taste responses could arise as a consequence of place cell remapping. However, we could not address this question in the current study, since rats did not explore the behavioral chamber in the absence of tastes, making it difficult to calculate place fields. We specifically chose a smaller open field environment for these experiments because it allowed adequate coverage of place fields as well as multiple taste stimuli, allowing us to study the real-time interaction between taste and place. Future studies that incorporate place-specific taste delivery in a larger or more structured environment, such as a linear track, will be able to explore whether taste experience can modify animals’ hippocampal representation of environments through rate or global remapping, as has been shown for other sensory modalities (Moita et al., 2004; Fyhn et al., 2007; Zhang and Manahan-Vaughan, 2015).

Whatever the relationship between spatial and gustatory firing, more hippocampal interneurons—by their very nature, non-place cells—responded to non-spatial stimuli than place cells (**Figure 2**). This result is consistent with studies that measured hippocampal responses to auditory (Moita et al., 2003) or olfactory (Deshmukh and Bhalla, 2003) stimuli. This finding may reflect the intrinsic properties of each cell type: place cells have lower mean firing rates and rarely respond outside of their place field, while interneurons exhibit high firing rates regardless of location (see **Figure 2**), making it much easier to obtain statistical significance in the latter. Another (though not mutually exclusive) possibility is that interneurons could be influencing place cell activity. While place cells’ responses to space have been well-characterized (O’Keefe and Nadel, 1978; Moser et al., 2008), recent work suggests that interneurons also contribute to hippocampal representations of space (Wilent and Nitz, 2007), and disinhibit place cell firing through location-specific decreases in activity (Hangya et al., 2010; Royer et al., 2012). Since interneurons’ projections are primarily internal to the hippocampal network (Freund and Buzsáki, 1996; Kullman, 2011), these responses may play a role in shaping activity within the network, such as place fields (**Figure 4**) or location-specific responses to tastes (**Figure 5**). However, how non-spatial information is transmitted within hippocampal microcircuits remains an open question, one that may be investigated in future studies by determining the effect of cell-type-specific inhibition on hippocampal taste responses.

There is strong evidence that the behavioral relevance of sensory stimuli within a task influences what proportion of hippocampal neurons respond to non-spatial cues. In our study, rats passively received tastes via IOC, a paradigm that requires no learning, other than associating tastes with a context for the first time. The total proportion of taste-responsive neurons in our study (~20%, **Figure 2**) is comparable to the proportion of tone-responsive cells (16%) found by one study analyzing the auditory evoked responses of hippocampal neurons (Moita et al., 2003); however, this proportion increased to 52% following auditory fear conditioning. Similarly, ~40% of hippocampal neurons exhibited non-spatial firing during an odor-guided non-match-to-sample task that required working memory of spatial and non-spatial factors (Wood et al., 1999). More pronounced changes in hippocampal responsiveness are observed when reward contingencies are entirely dependent on discriminating between non-spatial cues, such as in one study where rats learned to manipulate a joystick to modulate a tone within a target frequency range (Aronov et al., 2017). During this task, about 40% of hippocampal neurons responded to specific tone frequencies, compared to only 2% during the passive playback of tones. In our study, 8.3% (40/482) of CA1 neurons discriminated between tastes (**Figure 6A**), with an additional 11.6% (56/482) of cells responding nonspecifically to taste presence. These results suggest that the hippocampus forms a flexible map of spatial and non-spatial stimuli based on current behavioral demands. For the case of taste, gustatory responses in spatially tuned hippocampal neurons may allow animals to form value-related associations between tastes and contexts, aiding in the finding of food. Future work will assess how taste experience affects this ongoing mental map.

## Acknowledgements

This work was supported by a Sloan Research Fellowship in Neuroscience (Alfred P. Sloan Foundation) and Whitehall Foundation award to SPJ; NIH Grants R01 DC006666 and R01 DC007703 to DBK; Training grant T90 DA032435 to LEH.

